# Reconstructing Promoter Activity From Lux Bioluminescent Reporters

**DOI:** 10.1101/117093

**Authors:** Mudassar Iqbal, Neil Doherty, Anna M.L. Page, Saara N.A. Qazi, Ishan Ajmera, Peter A. Lund, Theodore Kypraios, David J. Scott, Philip J. Hill, Dov J. Stekel

## Abstract

The bacterial Lux system is used as a gene expression reporter. It is fast, sensitive and non-destructive, enabling high frequency measurements. Originally developed for bacterial cells, it has been adapted for eukaryotic cells, and can be used for whole cell biosensors, or in real time with live animals without the need for slaughter. However, correct interpretation of bioluminescent data is limited: the bioluminescence is different from gene expression because of nonlinear molecular and enzyme dynamics of the Lux system. We have developed a modelling approach that, for the first time, allows users of Lux assays to infer gene transcription levels from the light output. We show examples where a decrease in bioluminescence would be better interpreted as a switching off of the promoter, or where an increase in bioluminescence would be better interpreted as a longer period of gene expression. This approach could benefit all users of Lux technology.

## Introduction

The lux operon contains genes for the bacterial bioluminescent reaction [1, 2]: *luxA* and *luxB* encode the *α* and *β* subunits of the heterodimeric bacterial luciferase; *luxC* encodes a 54kDa fatty acid reductase; *luxD* encodes a 33kDa acyl transferase; and *luxE* encodes a 42kDa acylprotein synthetase. These genes, including their order (*luxCDABE*), are conserved in all lux systems of bioluminescent bacteria. An additional gene (*luxF*), with homology to *luxA* and *luxB,* is located between *luxB* and *luxE* in some species. The light emitting reaction, catalysed by the LuxAB complex, involves the oxidation of FMNH_2_ and the conversion of a long chain fatty aldehyde (tetradecanal *in vivo*) to its cognate acid, with the emission of blue-green light. LuxC, D and E together form the fatty acid reductase complex, involved in a series of reactions that recycle the fatty acid back to aldehyde. In *E. coli* and other species, Fre has been shown to be the enzyme responsible for flavin reduction back to FMNH_2_.

Gene expression can be measured by cloning a promoter of interest upstream of the lux operon, and interpreting the bioluminescence from bacteria containing such constructs as a measure of transcription [3, 4]. This provides a reporter that can measure gene expression at high frequency and with less background noise than other reporters, such as GFP [5, 4], and has found great value in both bacteria [6] and eukaryotes [7], with important recent applications in whole cell biosensors [8], live animal infection models [9, 10] and live tumour infection models [11].

However, this light is an integrated signal of transcription, mRNA halflife, translation and protein turn-over, the bioluminescence reaction kinetics and substrate availability and cycling. As a consequence, absolute transcription activity cannot be directly inferred from the data generated. This is a limitation in the current use of Lux technologies. For example, some studies using Lux reporters have observed fluctuations in light output [12, 13, 14, 15, 16], and it is not clear whether these truly reflect promoter activity, or are artefacts of the reporter system.

In this paper we show that detailed mathematical models for bioluminescence can be used to relate bioluminescence to promoter activity. We developed a new model that addresses limitations of previous work [17], whose structure and parameters are informed by new data and new mechanisms. We generated new enzymatic data both for the Fre reaction and for the LuxAB reaction, using *Photorhabdus luminescens* LuxAB that we have purified. An important part of our approach is to fit the models directly to the time series experimental data, which can be thought of as a (partial) factorial experiment with varying concentrations of both FMN and NADPH. This allows for the simultaneous inference of all parameters in complex kinetic models, which is an advantage over traditional chemical kinetics techniques that only use the maximal velocity.

## Results and Discussion

The luciferase reaction uses FMNH_2_ as an energy source. This is an energetically transient species with a short half-life. *In vivo* it is supplied to the luciferase complex for immediate consumption by the redox activity of the Fre enzyme, which converts NADPH and FMN to FMNH_2_ and NADP+. We measured the kinetics of the Fre reaction (Figure 1a) by recording the consumption of NADPH (as determined spectrophotometrically) at different starting concentrations of the FMN acceptor component (100*μ*M, 200*μ*M and 400*μ*M); in all cases, the initial NADPH concentration was 200*μ*M. The initial velocity of the reaction increased with increased FMN, leading to different kinetics in the three curves, with the steady state being reached more rapidly with increased FMN concentration.

**Figure 1:**
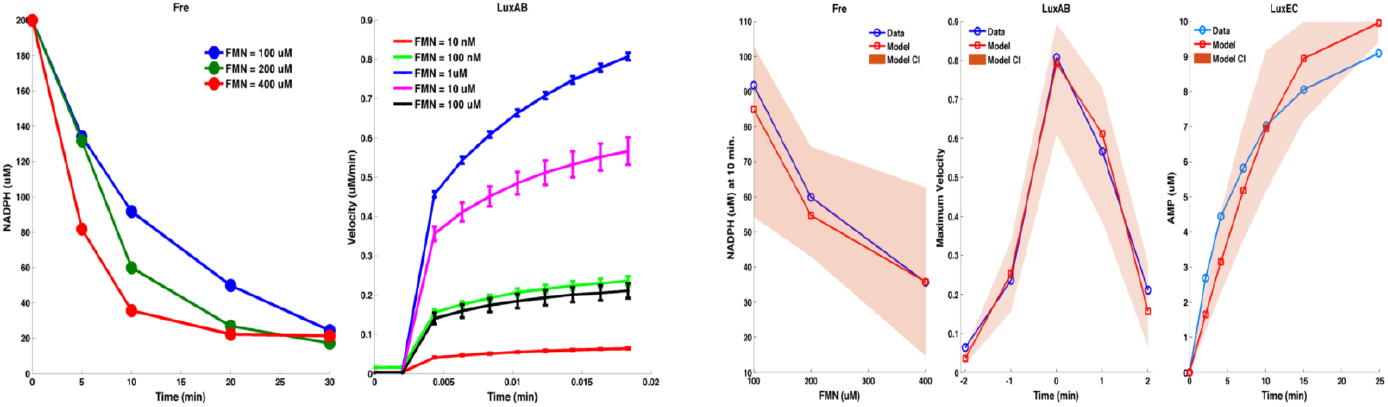
(a) New experimental data for Fre and LuxAB reactions. NADPH time course for three different concentration of FMN. The concentration values in the above data are obtained from the absorption measurements from the spectrophotometer. The conversion is carried out by relating the concentration (C) to the measured values (A) with the formula *A* = *K* * *C*, where *K* is the proportionality constant and is estimated using initial measurement(*A*_0_) and starting concentration (*C*_0_ = 200*μM*). The velocity of the reaction increases as FMN concentration is increased. Normalised (using max. velocity in the assay) LuxAB reactions velocity time series for different FMN concentrations. The velocity of the reaction increases from the lowest concentrations of FMN, is greatest for FMN concentrations of 1*μM*, and decreases for higher concentrations of FMN. This is best explained by product inhibition of the LuxAB reaction through competition between FMN and the substrate FMNH_2_. (b) Model fits for Fre, LuxAB, and LuxEC reactions. The model fits to the data are good, showing that kinetic parameters for the reaction rates can be inferred. Summarized data are displayed: for Fre - NADPH concentrations at t = 10 min; for LuxAB, the maximal veclocity for each FMN concentration; for LuxEC - only the AMP time-course data are shown. The full data fits for all three reactions are shown in supplementary figures. Total flavin = 88*uM*, *O*_2_ = 550*uM*, *NADPH* = 560*uM*, and *ATP* = 1310.

For the measurement of luciferase (LuxAB) kinetic reactions, we combined components of the coupled Fre-LuxAB reaction. We measured light output arising from different initial concentrations of FMN: 10nM, 100nM, 1*μ*M, 10*μ*M and 100*μ*M (Figure 1a). For all five conditions, there is an initial delay before light is produced, due to a two-step injection method. Following the lag, light is produced, initially rapidly, and then tailing off. A striking feature of these data is that maximum light production increases as FMN concentration is increased from 10nM to 1 *μ*M, but then decreases again as FMN concentration is increased further. The most likely explanation for this decrease is inhibition of the LuxAB reaction by its product, FMN, which would be competing with FMNH_2_ for binding to the LuxAB complex. It cannot be substrate inhibition of Fre by FMN as the kinetics for this reaction increase up to 400uM (Figure 1a). Product inhibition has not been previously reported and represents a newly discovered element of the Lux bioluminescent pathway. Structural studies have shown that both FMN and FMNH_2_ can form a complex with LuxAB [18], with only two residues that could act to discriminate between them (see PDB 3fgc), supporting the proposed product inhibition.

The new mathematical model includes three chemical reactions: the Fre reaction for flavin recycling; the LuxAB reaction for light production; and the LuxEC reaction for aldehyde recycling. It also contains a further equation to describe the turnover dynamics of the Lux proteins:

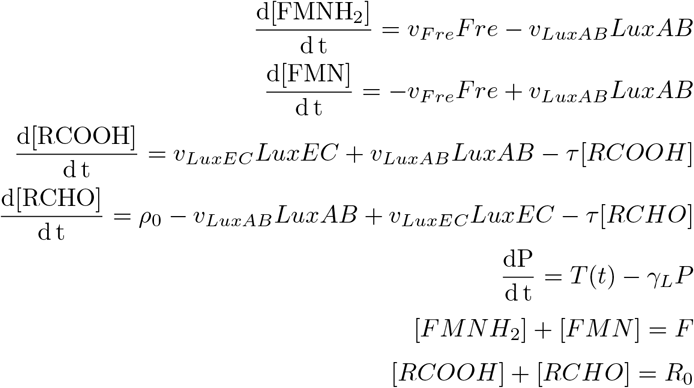

It is assumed that the action of LuxD counterbalances the loss of RCOOH and RCHO; thus *R*_0_ is constant and *ρ*_0_ = *τR*_0_. Control of the bioluminescent reactions is assumed to lie with the most rapidly turned-over protein. This is modelled by setting *P*(*t*) to represent all of the Lux proteins and using the same rate of synthesis and turn-over for all proteins. The protein production *T*(*t*) encompasses transcription, translation and mRNA degradation and is supplied as a model input; it is assumed that the *lux* mRNA is at quasi steady state. The value of the protein turn-over rate *γ_L_* has been inferred from Lux data as 0.378h^−1^ (Figure 2). Full details for the velocity equations for the terms *υ_Fre_*, *υ_LuxAB_* and *υ_LuxEC_* are provided in the Supplementary Materials. New detailed mechanisms have been defined using King and Altman's schematic method [19], with a modification to the *υ_LUXAB_* velocity term to include the impact of FMNH_2_ product inhibition.

**Figure 2:**
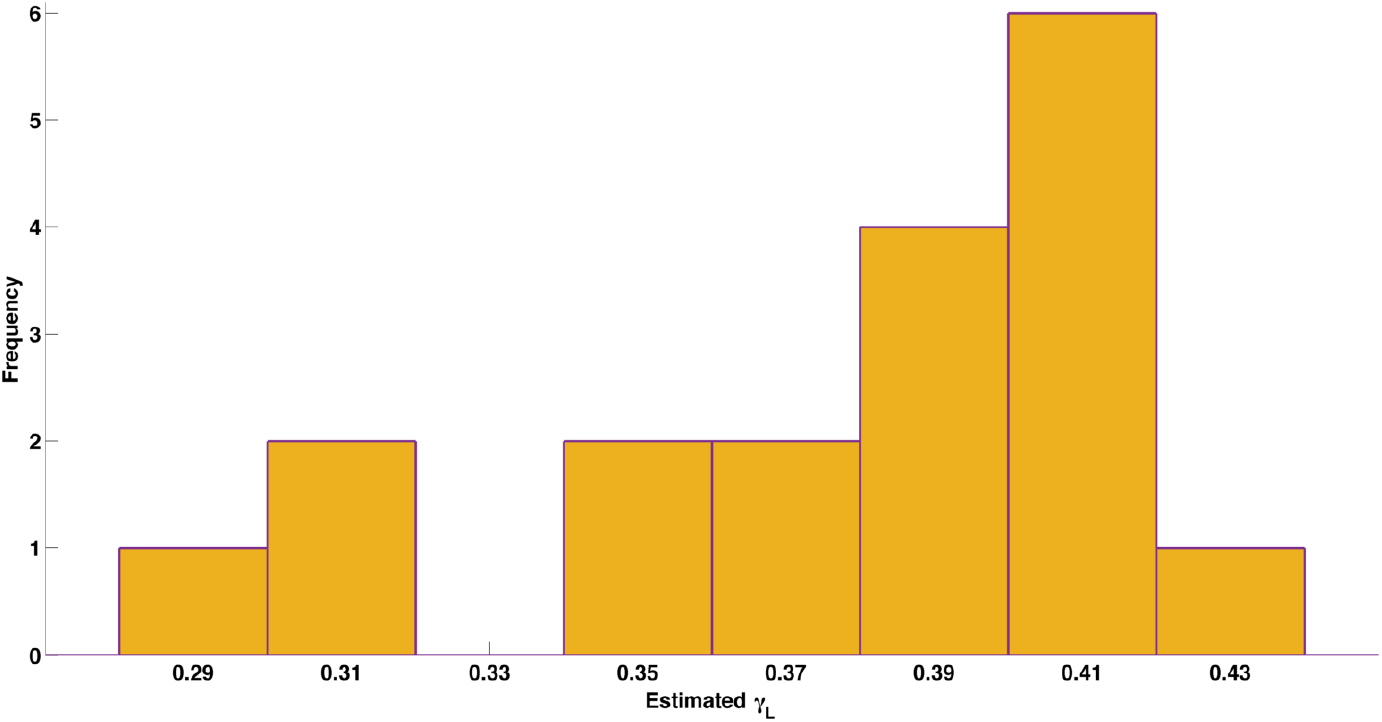
Histogram of inferred Lux protein turnover rates, showing low variability about a mean value of 0.378h^−1^.

We have carried out parameter inference for all three reactions: Fre, LuxAB and LuxEC, using a Markov Chain Monte Carlo (MCMC) approach, as in previous work [20, 21]. The results are posterior distributions for each of the parameters in the model, which describe not only the best fit values for the parameters, but also the degree of uncertainty of the parameter estimates. Because the Fre reaction is also used in the LuxAB experiment, inference for these parameters was carried out in two steps: first we fitted the experimental data from the Fre reaction to produce posterior distribution for these parameters; then we used these posterior distributions as prior distributions for the same parameters in the LuxAB reactions. For the LuxEC reactions, we used published data on steady-state measurements of NADPH, ATP and RCOOH, as well as time-course measurements of AMP formation [22, 23, 1], as shown in Figure 2 (B,F,G, H) of Welham et al. [17]. There is good concordance between the model and the data (Figure 1b), with the time course in particular showing substantial improvement (*R*^2^ = 0.762) over the model fit previously reported [17] (*R*^2^ = 0.396), consistent with improved model mechanism and parameter inference. Detailed fitted curves and MCMC diagnostics are shown in Supplementary Materials, and he inferred parameters are given in Supplementary Table 1.

A key finding of this analysis, of particular significance for inferring levels of gene expression with lux reporters, is that there is a nonlinear relationship between promoter activity and light output (Figure 3). Bioluminescent outputs display qualitative and quantitative behaviours different from the underlying gene expression, so direct interpretation of bioluminescent data can be misleading. Simulations of the model for a switching on of gene expression show that for low levels of expression, bioluminescence appears more gradually with a slight delay, while for higher levels of gene expression there is a transient peak of bioluminescence followed by lower steady state. Simulations for a transient pulse of gene expression also produce bioluminescent outputs that are different from the underlying gene expression. Bioluminescence decreases more slowly than gene expression, and, for higher levels of gene expression, bioluminescence can remain high long after gene expression has ceased. More detailed plots of all species in the reactions can be found in Supplementary Materials.

**Figure 3:**
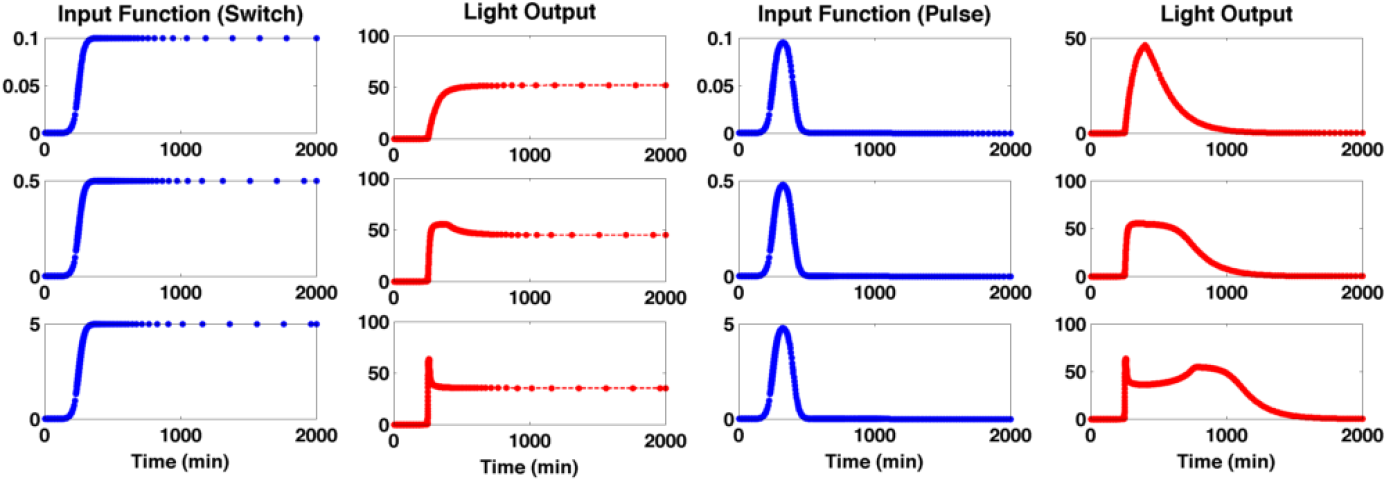
Relationship between promoter activity and light output Nonlinear relationship between promoter activity and light output for a synthetic pulse or switch of gene expression at different levels. The bioluminescence displays different qualitative and quantitative behaviours from the underlying gene expression. With the switch data, the bioluminescence has slower onset compared with gene expression, and, for high levels of gene expression, shows a transient pulse not present in the gene expression. With the pulse data, the bioluminescence shows much longer persistence than the underlying gene expression. These data show that bioluminescence alone could be a misleading measure of gene expression.

The model can be used to reverse engineer gene expression from bioluminescence, also using an MCMC approach. We have tested this approach on three datasets: the synthetic pulse data set, where the underlying gene expression to be inferred is supplied; data from the uhpT promoter in *S. aureus,* with relatively simple light profiles; and data from an acid stress experiment in *E. coli,* with more complicated bioluminescent output, for the promoter of the *safA-ydeO* operon in four strains following exposure to acid: wild-type, a *ydeO* knockout, a phoP knockout and a *ydeOphoP* double knock out; both YdeO and PhoP repress this promoter [15].

The reverse engineering of the synthetic pulse data show accurate reproduction of the supplied input gene expression profile, demonstrating that our method works correctly (Figure 4). For the uhpT data (Figure 5a), the inferred promoter activity suggests both earlier gene expression and more rapid switching off of expression than would be apparent from the light output. The most profound difference in interpretation of results based solely on levels of emitted light arise from the acid stress data (Figure 5b). While the inferred gene expression is less smooth than the bioluminescence, it highlights three behaviours not apparent in the bioluminescence itself. First, there is a clear, transient pulse of gene expression following acid stress, lasting only 20 minutes in the WT, 30 minutes in the single knockout strains, and 40 minutes in the double knock-out. In the WT and *ydeO* mutant, gene expression is completely switched off after this pulse, which cannot be seen from the bioluminescence, while in the *phoP* and double mutants, it is not completely switched off. Second, the increase in gene expression in the mutants relative to the WT is much lower than indicated by the bioluminescence, with the increased bioluminescence reflecting increased duration of gene expression as much as increased level. Third, the inferred gene expression appears to show pulses. These reflect the experimental protocol, in which plates were moved between the luminometer and the spectrophotometer every 15 minutes, disturbing the cells. The pulses suggest that the protocol had a more profound impact on cell activity than would be apparent from the light. The peaks of these pulses are more likely to represent gene expression level, indicative of a long term steady state gene expression, also not apparent from the bioluminescence. While the transient gene expression of the *safA-ydeO* promoter is to be expected in the WT and two single mutant strains, the reason for its transience in the double mutant, i.e. in the absence of the two known down-regulators of the promoter, is not known. The promoter is activated by the EvgA response regulator following a drop in pH, and the kinetics of turnover and dephosphorylation of this activator are unknown. They may explain the transient activation of the promoter, or there may be other feedback systems operating. Other known acid responsive regulators such as GadE, GadX and GadW are not responsible, as their deletion has no affect on promoter induction kinetics [15].

**Figure 4:**
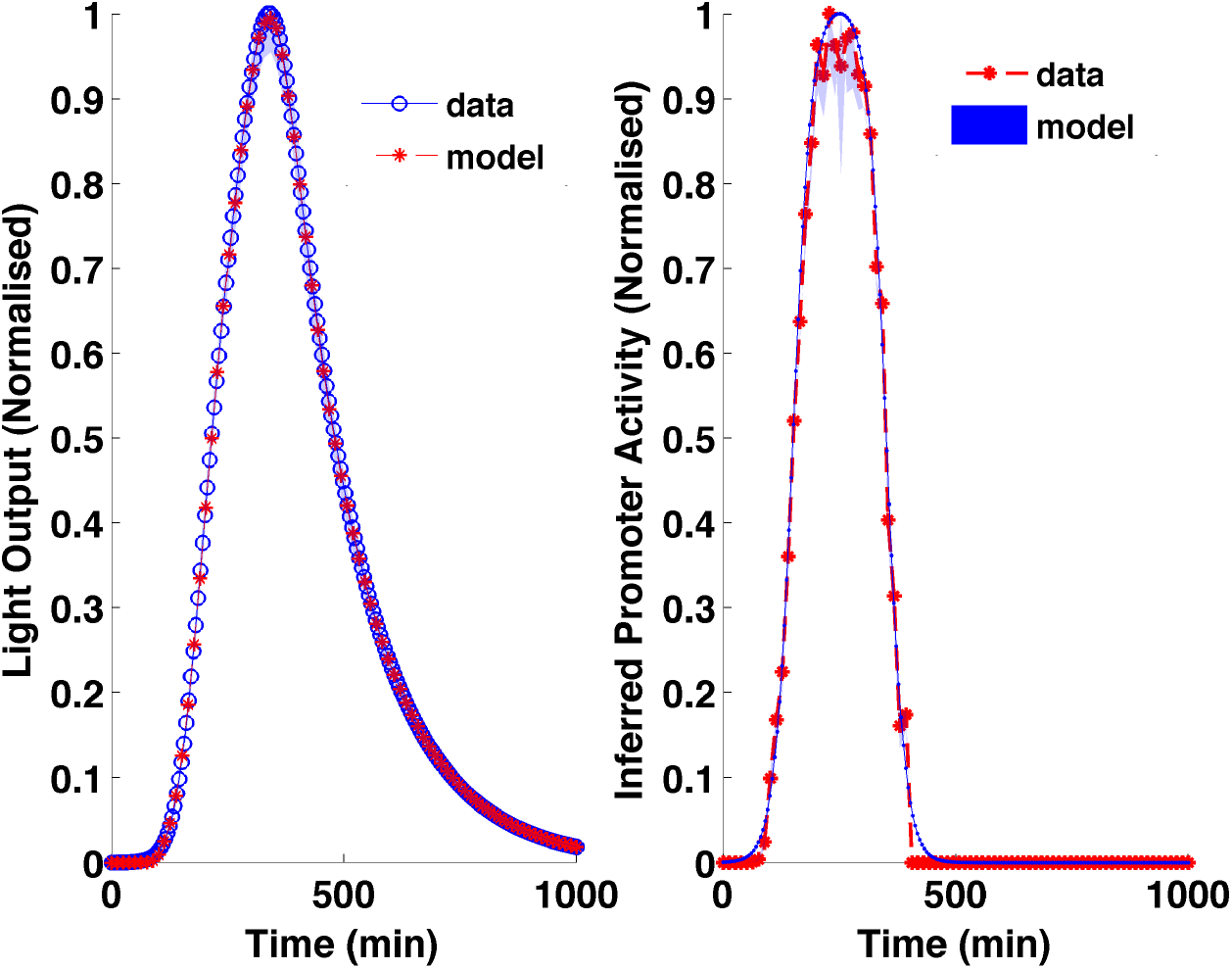
Reverse engineered promoter activity from light output from a simulated transient pulse experiment showing effective and accurate recovery of the known gene expression profile.

**Figure 5:**
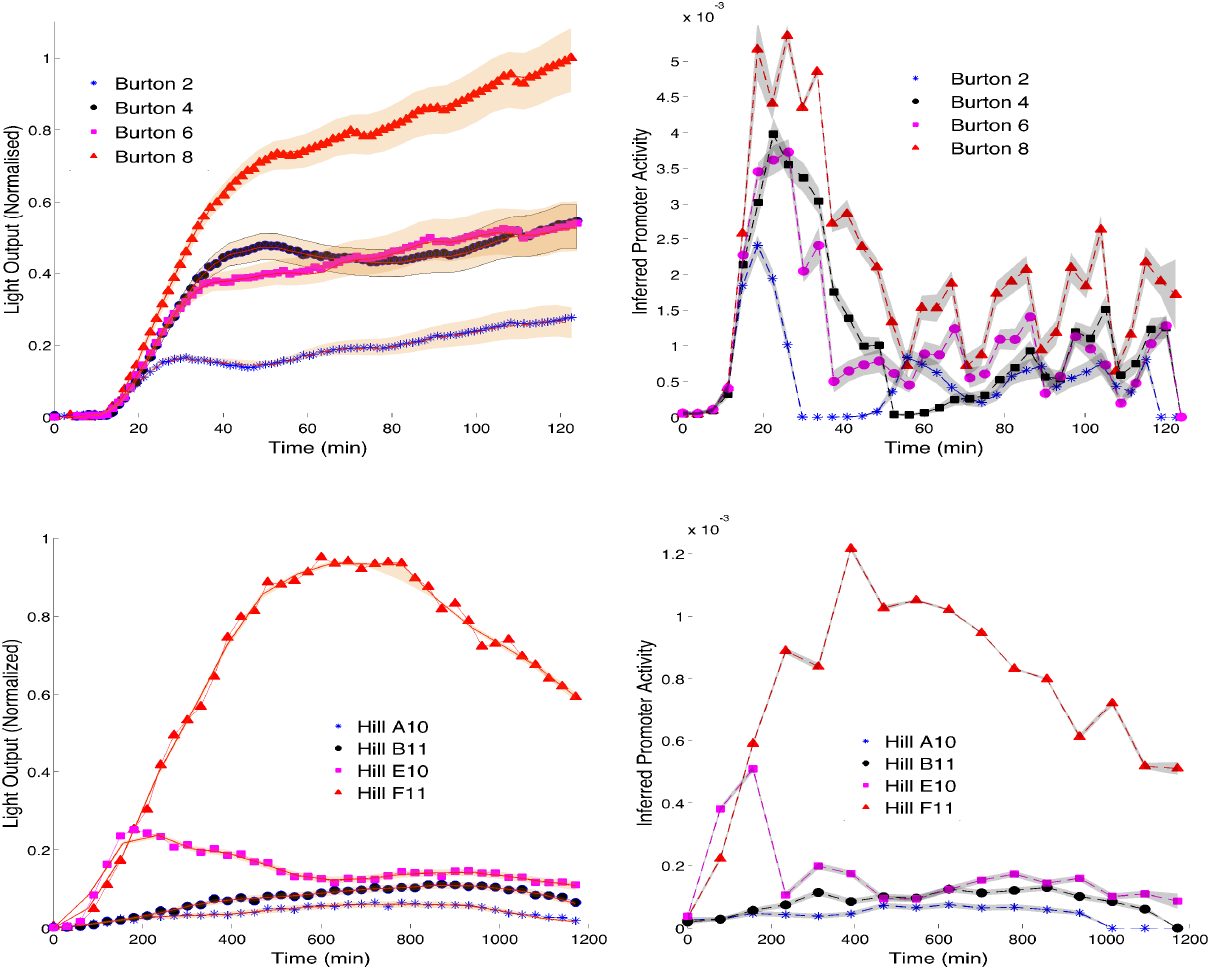
(a) Reverse engineered gene expression from data from the *uhpT* promoter in *S. aureus* showing that the peak of gene expression occurs earlier than the light output, and that the duration of gene expression is shorter than would appear from the light output. (b) Reverse engineered gene expression from the *safA-ydeO* promoter in *E. coli.* The pattern of gene expression is very different from the light output pattern. In particular, the increased bioluminescence is partly explained by greater duration of gene expression not just level of gene expression; gene expression in the WT and *yodel* mutant appears to be switched off after induction, which is not apparent in the bioluminescence; and the inferred expression shows pulses which are an artefact of the experimental arrangement, in which plates were moved every 15 minutes between a spectrophotometer and a luminometer, demonstrating that the agitation has an impact upon the cells.

In conclusion, we have developed a new mathematical model to relate gene expression to light output in Lux promoter assays, that includes newly discovered experimental evidence for product inhibition of the LuxAB reaction by FMNH_2_. The model shows a nonlinear relationship between gene expression and light output. We have used the model to provide a method to infer gene expression from light output that can be generally applied to experiments using Lux reporter assays. We anticipate that this approach could have valuable applications in inferring gene expression levels in a wide range of biological systems where lux reporters can be employed, including both *in vitro* experiments, and to track gene expression in animal models of bacterial infection. Program code to undertake reverse engineering of promoter activity are provided as Matlab and R functions in supplementary files.

## Methods

### Chemicals, media, cloning, bacterial strains

All chemicals were purchased from Sigma-Aldrich unless indicated otherwise, and were of ultrapure quality. Routine cloning steps were carried out using standard molecular biological protocols. Primary clones were selected in LB with Mach I Electrocompetent or chemically *E. coli* obtained from Thermo-Fisher Scientific. Protein expression was carried out using the *E. coli* strain BL21 (DE3) which had been previously transformed with the arabinose inducible plasmid pGRO7, encoding the GroES-GroEL chaperone complex (Takara Inc.). Autoinduction medium was used for the protein purification steps (Overnight Express Autoinduction system 1—Novagen). See below for further details of the luciferase purification strategy. Antibiotic selection was carried out at the following concentrations: Chloramphenicol - 20*μ*gml^−1^, Carbenicillin - 100*μ*gml^−1^, Kanamycin - 50*μ*gml^−1^.

### Measurement of Fre activity

The commercially available Fre/NAD(P)H:FMN-Oxidoreductase of *P. luminescens* was obtained from Roche Diagnostics. The activity of the enzyme was determined using a continuous spectrophotometric rate determination method as follows: NADPH (200*μ*M), FMN (100-400*μ*M) and Fre (5*μ*l of a 0.2 unit ml^−1^ preparation per ml final reaction mix) was prepared in 50mM potassium phosphate buffer, pH 7.0. Replicate 1ml samples were prepared in sterile cuvettes, which were mixed by inversion immediately after addition of the enzyme. Fre activity was observed as a function of the loss of NADPH, measured by reduction of its characteristic absorbance value at 340nm. Experiments were conducted over a period of 30 minutes at 22° C. All measurements were performed in triplicate.

### Luciferase purification

A co-expression approach was undertaken to purify nascent LuxAB directly from *E. coli* BL21 (DE3). The luciferase genes were amplified from the plasmid pSL1190::luxA2E bearing the *P. luminescens* luciferase operon using the high fidelity polymerase KOD (Merck-Millipore Corp.). *luxA* (Genbank ID AAA276619.1) was amplified using the primer pair: luxAfw 5’-AGCACGCATATGGCGAAATTTGGAAACTTTTTGCTTACA-3’ / luxArv 5’-CCGTCGCTCGAGTTAATATAATAGCGAACGTTG-3’. *luxB* (Genbank ID AAA27620.1) was amplified using the primer pair: luxBfw 5’-GAGCACGCATATGGCCAAATTTGGATTGTTCTTCC-3’ / luxBrv 5’-CCGTCGCTCGAGTTAGGTATATTTCATGTGGTACTTC-3’. *luxA* and *luxB* were cloned into pET21b and pET28b respectively, using a *XhoI*/*Nco*I digest approach. A stop codon was placed in the *luxArv* primer, leaving an N-terminally tagged LuxB and an untagged LuxA in the final expression system. These plasmids were co-transformed into a version of the expression strain which had previously been transformed with the plasmid pGro7, encoding the GroES/GroEL chaperone system. We found that this improved yield of soluble LuxAB. Following overnight growth of the co-expression strain in autoinduction medium containing the appropriate antibiotics and arabinose at 1mgml^−1^ (37°C with shaking), cells were harvested by centrifugation and lysed with the Bugbuster cell lysis reagent with added benzonase and lysozyme according to the manufacturer's instructions (Merck-Millipore Corp.). Clarified lysate was diluted five-fold in wash/bind buffer (20mM tris, 200mM NaCl, pH7.4) containing protease inhibitors (Protease Inhibitor Cocktail—Sigma-Aldrich co.). The resulting preparation was applied to equilibrated Ni^2+^ sepharose columns (HiTrap - GE Healthcare Life Sciences), washed with 100 column volumes wash/bind buffer, and luciferase was eluted in 5 × 1ml fractions of elution buffer (wash/bind buffer containing 500mM imidazole). Active fractions were assessed using the reconstituted luciferase assay (see section M4) using a Biospacelab Photon Imager, and combined. The N-terminal his-tag was cleaved from LuxB using 3 units ml^−1^ cleavage-grade thrombin according to the supplier's instructions (Novagen). Cleaved luciferase was buffer exchanged and purified from the released tag and other low molecular weight contaminants such as imidazole using 10kDa nominal molecular weight cut-off centrifugal filter units (Microcon YM10—Sigma-Aldrich Co.). Protein concentration was determined by Bradford assay using a Nanodrop low sample volume UV-Vis Spectrophotometer (Thermo Scientific) using a BSA standard curve.

### Reconstitution of coupled luciferase assay and kinetic measurements

Purified luciferase and commercially prepared Fre were combined to form the coupled reaction complex as follows: Reactions were typically carried out in a final volume of 100*μ*l containing purified luciferase (typically in the range 1.5-15*μ*gml^−1^), NADPH (200*μ*M), FMN (tested over the range 10nM-100pμ M in this study), Fre (0.2 units ml^−1^) and decanal (0.02%) in 50mM potassium phosphate buffer, pH 7.0. Reactions were carried out at 22°C and monitored in either a Biospacelab Photon Imager or Tecan Genios Pro multimode microplate reader. The kinetic assays were carried out using a Tecan Genios pro fitted with injector capacity as follows; all components other than decanal were combined in a final volume of 50*μ*l per well in flat bottomed microtiter plates suitable for bioluminescence measurements. The reactions were initiated by injection of a further 50pμ l phosphate buffer containing 0.04% decanal. Reactions were monitored for 1100 milliseconds. All measurements were performed in triplicate.

### Estimation of Lux Turnover Rates

The promoter of the Universal hexose transporter (*uhpT*) gene was amplified from *S. aureus* and introduced into pUNK1dest along with the Gram-positive GFP-*lux*ABCDE operon and the *rrn*BT1T2 terminator [24] using a Multisite Gateway LR plus reaction. Transformants were selected on erythromycin and screened for expression of the reporter. This PuhpT-reporter vector was designated pSB3009. *S. aureus* RN4220 [pSB3009] overnight cultures were grown aerobically at 37°C in in Tris Minimal Succinate medium (TMS, [25]) supplemented with Erythromycin (5*μ*gml^−1^) for plasmid maintenance Bacterial pellets were washed once in TMS without sodium succinate (TM) and resuspended in an equal volume of TM. These were diluted 1/50 into fresh TM supplemented with filter sterilized sugars (Glucose or glucose-6-phosphate) supplemented with Erythromycin (5*μ*gml^−1^).

For growth and reporter gene measurements, replicate samples (200*μ*l) were placed into the wells of a 96-clear-bottom microtiter plate (Porvair) and incubated at 37°C in a Tecan Genesis Pro microplate reader. Optical density (600nm), fluorescence (485ex/510em) and bioluminescence (RLU) readings were taken at 30 min periods over the course of the experiment.

From the data generated from these experiments, we identified the curves where RLU (light read-outs, arbitrary units) decreases while cells are still in exponential phase (corresponding OD measurements). We estimate the turnover rate for each such curve (total 18 curves used) by fitting a linear line to the log transformed data. The histogram of estimated values are shown in Figure 1d. The linear fits to the log-transformed data for all the 18 curves is shown in Supplementary Material.

### Parameter Inference

In order to estimate the kinetic parameters for all three reactions, we used a similar Bayesian approach based on Markov Chain Monte Carlo (MCMC), in which a sub-model relevant to each reaction is used along-with corresponding experimental data. We used an adaptive version of Metropolis-Hasting MCMC algorithm with global scaling [26] in order to iteratively sample from the posterior distribution of the kinetic parameters. Our choice of priors and likelihood function is described below:

Unless otherwise specified, we used un-informative exponential priors (*λ* = 100) for kinetic parameters, while more informative *gamma* priors (*Gam*(0.9, 0.1)) for degradation rates. This reflects our *a priori* knowledge that the kinetic parameters (ratio of kinetic rate constants) are all positive real numbers. For those parameters which are common to the inference in both the Fre and LuxAB models, prior estimates for the LuxAB inference were generated by fitting exponential distributions to the posterior estimates derived from the the Fre inference.

We define the likelihood of the parameters, for any of the models using in equation 1, assuming homogeneous Gaussian noise. Given the current set of parameter values (*θ*), we simulate the species of interest (stored in vector *Y′*), and use the corresponding experimental data in vector *Y* to give the likelihood function:

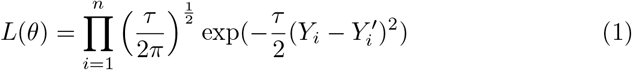

A separate Gibbs step is introduced for the sampling of noise precision *τ*, in case of Fre and LuxEC inference, while for LuxAB data, we estimated the noise variance from the replicates. Details on the derivation of the Gibbs step is provided in supplementary methods.

### Promoter Inference

A Monte Carlo approach was used to infer promoter activity from light readout. The promoter input function is modelled as a series of K heights at fixed positions. A Martingale prior distribution is used [27], so that the prior distribution for each height at point *n* is an exponential distribution with mean value equal to the current height at the previous point *n* − 1. At each step, a point is chosen at random, and a new height is proposed, as described in Green 1995[28]. The likelihood function uses a Gaussian error model. For all promoter inference, light output curves for a whole experiment must be normalized to the highest light value found in that experiment.

## Acknowledgments

This work was funded by BBSRC grant BB/I001875/1. We thank Tania Pere-hinec for technical support.

